# The regulatory network of the White Collar complex during early mushroom development in *Schizophyllum commune*

**DOI:** 10.1101/2024.02.13.580115

**Authors:** Peter Jan Vonk, Marieke J. P. van der Poel, Zoé E. Niemeijer, Robin A. Ohm

## Abstract

Blue light is an important signal for fungal development and is detected by the White Collar complex, which consists of WC-1 and WC-2. Most of our knowledge on this complex is derived from the ascomycete *Neurospora crassa*, where both WC-1 and WC-2 contain GATA zinc-finger transcription factor domains. In basidiomycetes, WC-1 is truncated and does not contain a transcription factor domain, but both WC-1 and WC-2 are still important for development . In the model mushroom *Schizophyllum commune*, we show that dimerization of WC-1 and WC-2 happens independent of light, but that induction by light is required for promoter binding by the White Collar complex. Furthermore, the White Collar complex is a promoter of transcription, but binding of the complex alone is not always sufficient to initiate transcription. For its function, the White Collar complex associates directly with the promoters of structural genes involved in mushroom development, like hydrophobins, but also promotes the expression of other transcription factors that play a role in mushroom development.

## Introduction

Light is an important signal in many filamentous fungi, affecting a large variety of processes, including metabolism, stress and development. While photoreceptors for red, green and blue light have been identified in fungi, the detection of blue light is the most ubiquitous and best-studied light signal in fungi. This is primarily due to the study of photobiology in the model organism *Neurospora crassa* (Chen et al., 2010; Dunlap and Loros, 2005). In this organism, the detection of blue light entrains the circadian clock, which affects many processes, including sexual and asexual development, sporulation and growth rate (Dunlap and Loros, 2017). Blue light is detected by the White Collar complex (WCC) a heterodimer consisting of WC-1 and WC-2 (Ballario et al., 1996; Linden and Macino, 1997). The function of these proteins in light sensing has previously been reviewed in detail (Corrochano, 2019). Both WC-1 and WC-2 contain a GATA zinc-finger transcription factor domain for DNA-binding. In addition, WC-1 contains a light-oxygen-voltage (LOV) domain that can bind a flavin chromophore and is responsible for the structural changes of the WCC upon the detection of blue light (He et al., 2002). Two activated LOV-domains can interact, resulting in a WCC dimer (Malzahn et al., 2010). Monomers of the WCC are able to bind the promoters of light-regulated genes, but dimerization is required for the activation of transcription. The binding of WCC dimers results in the opening of the chromatin by displacement of nucleosomes, and the recruitment of the histone acetyltransferase NGF-1 that acetylates histone H3 on lysine 14 (H3K14ac), promoting transcription (Grimaldi et al., 2006; Sancar et al., 2015). The process is negatively regulated by phosphorylation of the WCC, which prevents it from binding the promoter of *frq*, which encodes the clock-protein FRQ (Schafmeier et al., 2005). Furthermore, the WCC promotes transcription of *vvd*, which encodes the photoreceptor VVD. This protein disrupts dimerization of the WCC by binding to the WC-1 LOV-domain upon activation by light, acting as an inhibitor of the WCC.

Orthologs of WC-1 and WC-2 are found across the fungal kingdom, including mushroom-forming basidiomycetes (Idnurm et al., 2010; Ohm et al., 2013; Todd et al., 2014). This is expected, as blue light is an essential signal for the development of fruiting bodies in many species, including *Coprinopsis cinerea, Flammulina filiformis* and *Schizophyllum commune* (Kamada et al., 2010; Li et al., 2023; Ohm et al., 2013). In *S. commune*, deletion of either *wc-1* or *wc-2* prevents mushroom development completely, for which light is an essential signal (Ohm et al., 2013). In addition, the mycelium becomes more sensitive to UV-light. This is presumably due to a lower expression of ferrochelatase and DNA photolyase, which are important for preventing light-induced DNA damage. The interaction of WC-1 and WC-2 was shown for the first time in a basidiomycete by a yeast two-hybrid assay in *Pleurotus ostreatus* (Qi et al., 2020). Despite these similarities in function between the WCC in Ascomycota and Basidiomycota, there are also notable differences in the structure of the WCC between these phyla. In basidiomycetes, the WC-1 protein is truncated and lacks a GATA zinc-finger transcription factor domain. Furthermore, no ortholog of VVD has been identified in any basidiomycete (Idnurm et al., 2010). Finally, in the basidiomycete *Cryptococcus neoformans*, the number of light-regulated genes is much lower than in *N. crassa* (Chen et al., 2009; Idnurm and Heitman, 2010). This raises the question whether the mechanism and role of the WCC is conserved between different fungi.

Here we use the model mushroom-forming basidiomycete *S. commune* to study the activity of WC-2 in the presence and absence of light during mushroom development by a combination of transcription factor ChIP-Seq and RNA-Seq. This reveals the regulatory network of the WCC for the first time in a mushroom-forming fungus and shows the similarities and differences in light-sensing between ascomycetes and basidiomycetes.

## Results

### Yeast two-hybrid shows interaction of WC-1 and WC-2 independent of stimulation by light

To examine if *S. commune* WC-1 and WC-2 form a heterodimerize, the coding sequences of *wc-1* and *wc-2* were fused to the *GAL4* activating domain and *GAL4* binding domain, respectively. These plasmids were used to transform the *Saccharomyces cerevisiae* strains Y8800 and Y8930, respectively, to create two compatible fusion strains for yeast two-hybrid. Interaction assays revealed that WC-1 and WC-2 indeed interact in a Y2H assay (Figure 1). To determine the influence of light on this process, the experiment was repeated in the absence of blue light. Interaction still occurred, indicating that the formation of the WCC is independent of blue light.

**Figure 1:**
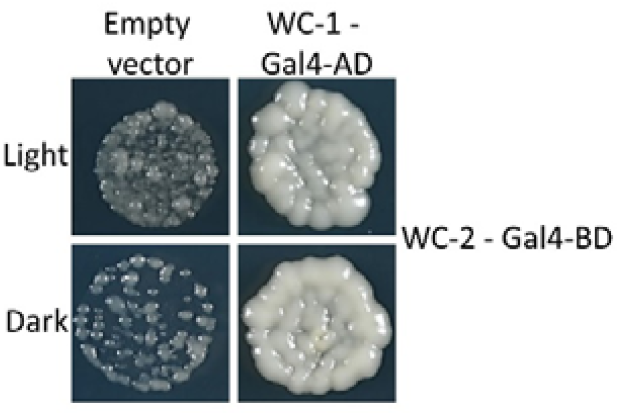
WC-1 and WC-2 interact in both light and darkness. Yeast two-hybrid of *wc-1* fused to the Gal4-activating domain (AD) and *wc-2* fused to the Gal4-binding domain (BD) in the light and the dark. The yeast is auxotrophic for histidine due to replacement of the *his3* promoter with the *gal4* promoter. Cultures are grown on yeast drop-out medium lacking adenine. Interaction of WC-1 and WC-2 results in the co-localization of the Gal4 AD and BD, promoting his3 transcription and complementing the *histidine* auxotrophy. In both the light and the dark adenine auxotrophy is restored when yeast is transformed with both WC-1 and WC-2 fusion proteins, while an empty vector with only the Gal4-activating domain does not complement the auxotrophy. Additional controls are depicted in Supplementary Figure 2.

### The WCC only associated with the promoters of genes in the presence of light

To study the association of the WCC with the promoters of light-activated genes, we employed transcription factor ChIP-Seq of WC-2 in both the light and the dark. Cultures were harvested right before the first signs of light-dependent fruiting became apparent (90 hours after inoculation). First, a Δ*wc-2*Δ*ku80* strain of *S. commune* was transformed with a plasmid encoding the full length *wc-2* gene including a C-terminal HA-tag. After crossing, this resulted in two Δ*wc-2* :: *wc-2-HA* strains with compatible mating types. When crossed, these strains complemented the Δ*wc-2* phenotype and were able to fruit (Figure 2). The presence of the protein could be detected by western blot (Supplementary Figure 1). The protein was then used for a pulldown of the chromatin of 90 hour old wild type and Δ*wc-2* :: *wc-2-HA* dikaryons with an anti-HA antibody. After purification and sequencing of DNA isolated during the ChIP, the binding sites of WC-2 could be identified. The vast majority of sites where WC-2 is associated with the chromatin were only occupied when the colony was illuminated. In the light a total of 569 binding sites were detected, while in the dark only two binding sites were identified (Supplementary Table 1). This indicates that light is required for the vast majority of DNA-binding activity of WC-2.

**Figure 2:**
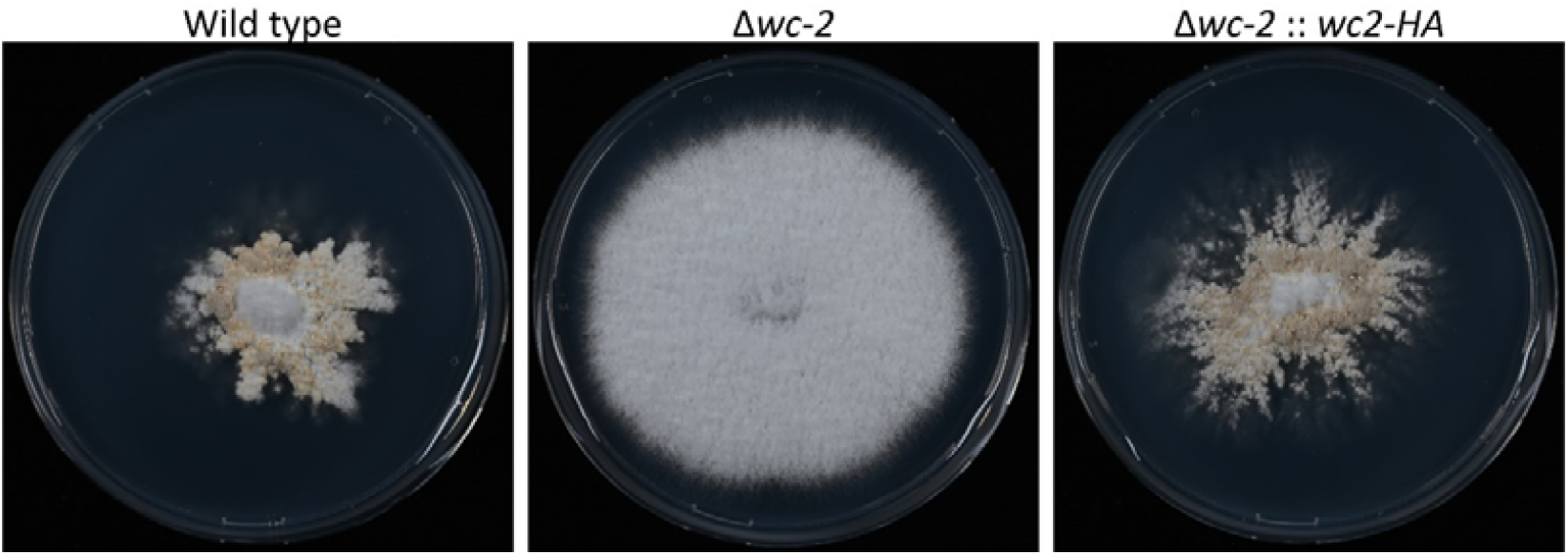
WC2-HA fully complements the *Δwc-2* phenotype. Cultures of wild type, Δ*wc-2* and Δ*wc-2* :: *wc-2-HA* dikaryons were grown for 7 days in the light. While a Δ*wc-2* dikaryon does not show any mushroom development for the wild type and Δ*wc-2* :: *wc-2-HA* dikaryon develop mature mushrooms.

The binding sites of WC-2 are enriched in the predicted promoter regions, upstream of the start codon of genes (Figure 3A). A binding site was associated with a gene if the tip of the peak was within 1000 bp of the translation start site. This resulted in the 569 peaks being associated with 549 genes (Supplementary Table 2). The promoters of several previously characterized genes were occupied by WC-2, including the hydrophobins *sc1, sc4, hyd1* and *hyd7*, and transcription factors *bzt1, fst3, hom2, pri2, tea1* and *zfc7*. All of these genes, except *pri2*, have previously been associated with various stages of mushroom development. In total, the promoters of 17 transcription factors were associated with WC-2 binding, indicating that WC-2 has a role in a larger regulatory network (Table 1). It was previously proposed that DNA photolyase *cry1* (protein ID Schco3|2621816) and ferrochelatase *fer1* (protein ID Schco3|2634259) were direct targets of the WCC (Ohm et al., 2013). Indeed, both genes are directly associated with WC-2 binding (Figure 3B, C).

**Table 1:**
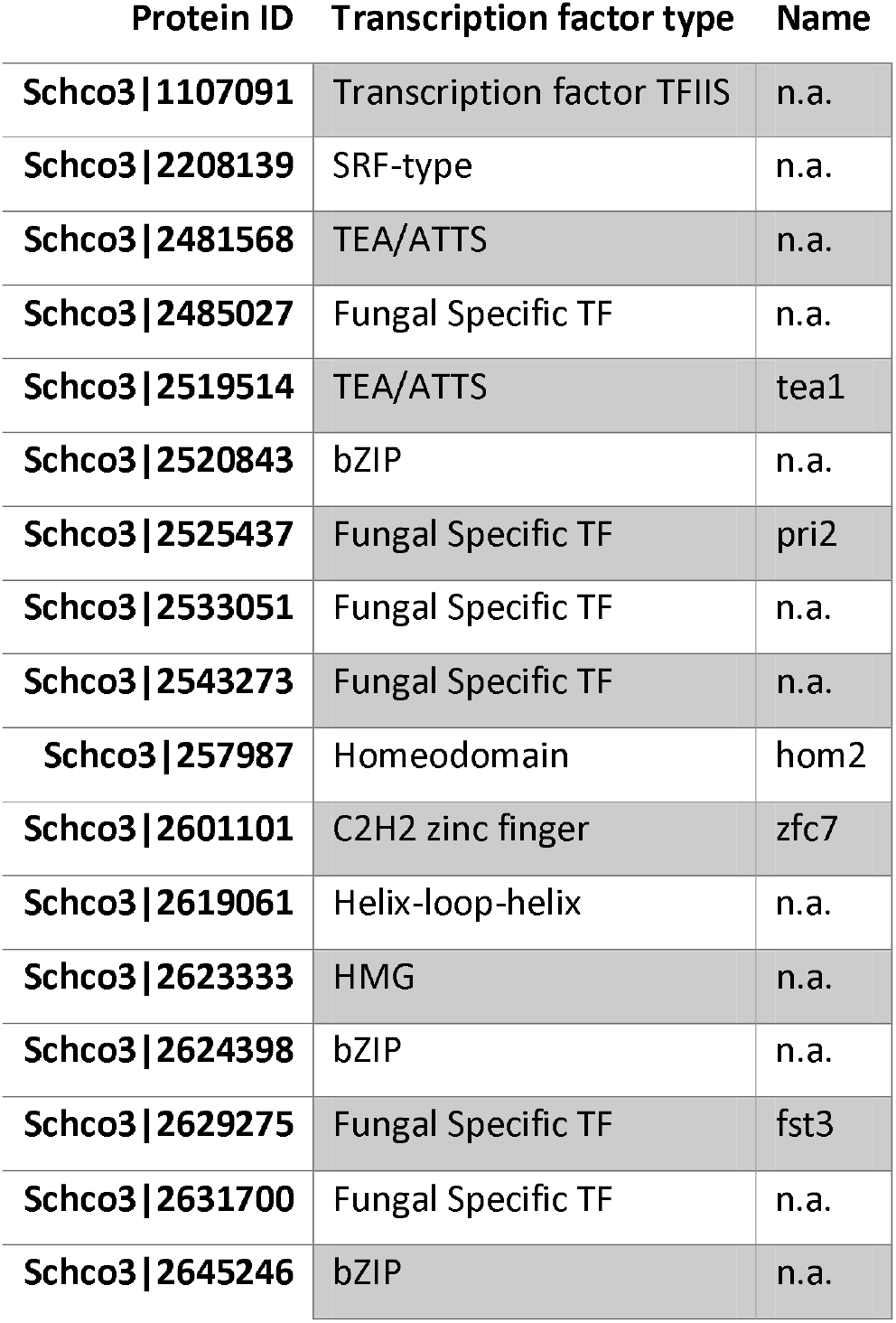
Transcription factors that are associated with a WC-2 binding site in the light. The predicted DNA-binding domain of each transcription factor is listed. Four of the transcription factors associated with a WC-2 binding site have previously been characterized: Fst3, Hom2, Pri2 and Zfc7 (De Jong et al., 2010; Ohm et al., 2011, 2010; Vonk and Ohm, 2021).

**Figure 3:**
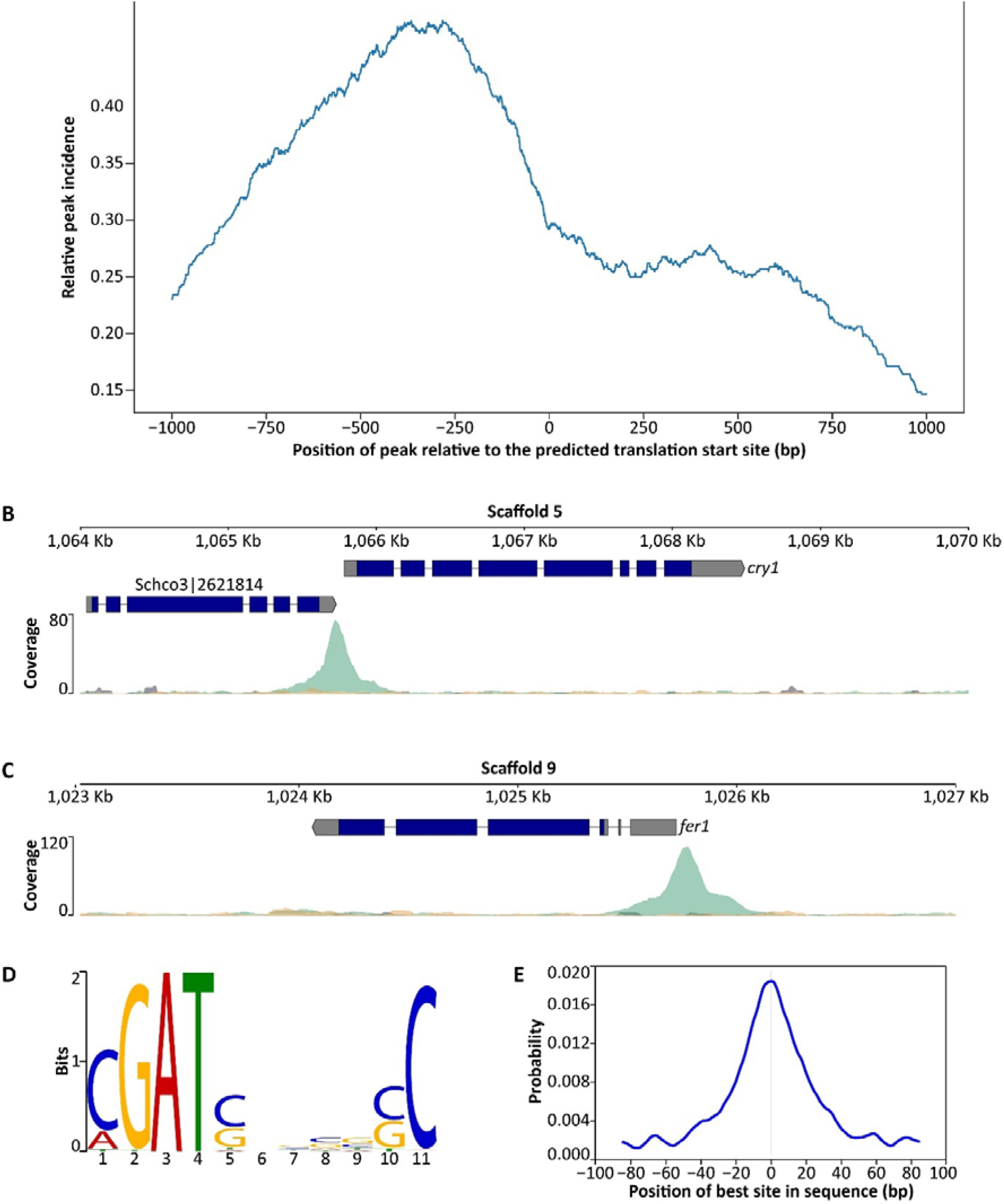
WC-2 activates transcription by binding the promoters of light-regulated genes that contain the motif CGATSNNNNSC. A: The relative occurrence of binding sites in the 1 kb upstream to 1 kb downstream of the translation start site (TSS). WC-2 preferentially binds upstream of the TSS of genes in the promoter region. B: ChIP-Seq peak of WC-2 in the promoter of the DNA photolyase *cry1* (protein ID Schco3|2621816). WC-2 binds upstream of the cry1 TSS. C: ChIP-Seq peak of WC-2 in the promoter of the ferrochelatase *fer1* (protein ID Schco3|2634259). WC-2 binds upstream of the fer1 TSS. D: The predicted consensus motif recognized by WC-2. E: The predicted motif is enriched in the center of WC-2 peaks, indicating that the motif is important in the interaction of WC-2 and chromatin.

To identify a putative binding motif of WC-2, the 200 bp around the tip of each peak was used for a motif enrichment analysis and novel motif discovery. The consensus motif CGATSNNNNSC was enriched in the sequences, occurring in 305 of the 565 sequences (Figure 3D). Moreover, the motif was centered around the tip of the peak associated with a predicted WC-2 binding sites, as is expected for a binding site (Figure 3E).

### RNA-Seq reveals that WC-2 promotes the expression of genes

Light is necessary for the association of WC-2 with the promoter sites of genes in *S. commune*. However, it is unclear if WC-2 promotes or inhibits transcription in *S. commune* and if light is sufficient for this activity. To examine the effect of WC-2 on transcription, we determined the differentially expressed genes (DEGs) between the light and dark in the WT and a Δ*wc-2* dikaryon at the same timepoint as in the ChIP-Seq analysis. A total of 271 genes were differentially expressed in any condition (minimum 4-fold difference in a condition compared to any other condition), the majority of which (195 genes) were differentially regulated in the WT between light and dark conditions (Figure 4, Table 2, Supplementary Table 3 & 4). In contrast, in the Δ*wc-2* dikaryon, only two genes were differentially regulated between light and dark. This shows that the Δ*wc-2* dikaryon does not respond to light and is effectively blind. Indeed, while there are 218 DEGs between the WT and Δ*wc-2* in the light, only 9 DEGs were identified in the WT in the dark compared to Δ*wc-2* in the light (Table 2).

**Table 2:**
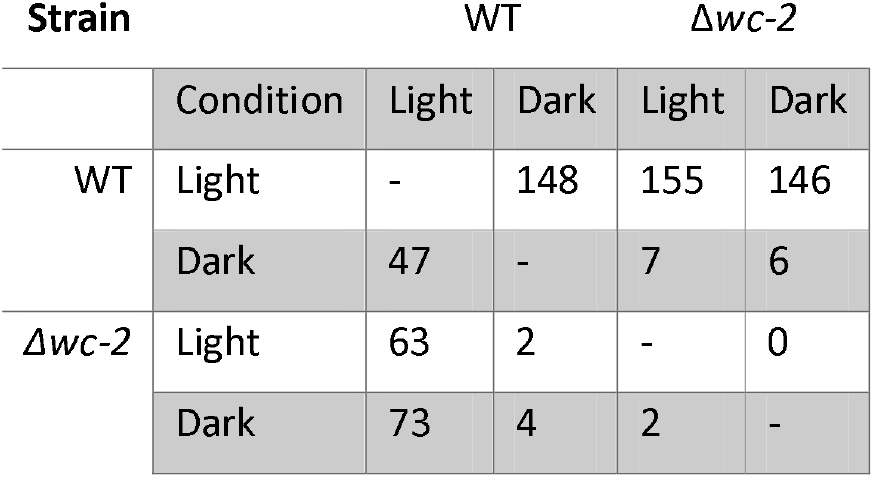
Upregulated genes between wild type and Δ*wc-2* dikaryons grown in the light and dark for 90 hours. Numbers indicate the number of upregulated genes in the horizontal condition compared to the vertical condition.

**Figure 4:**
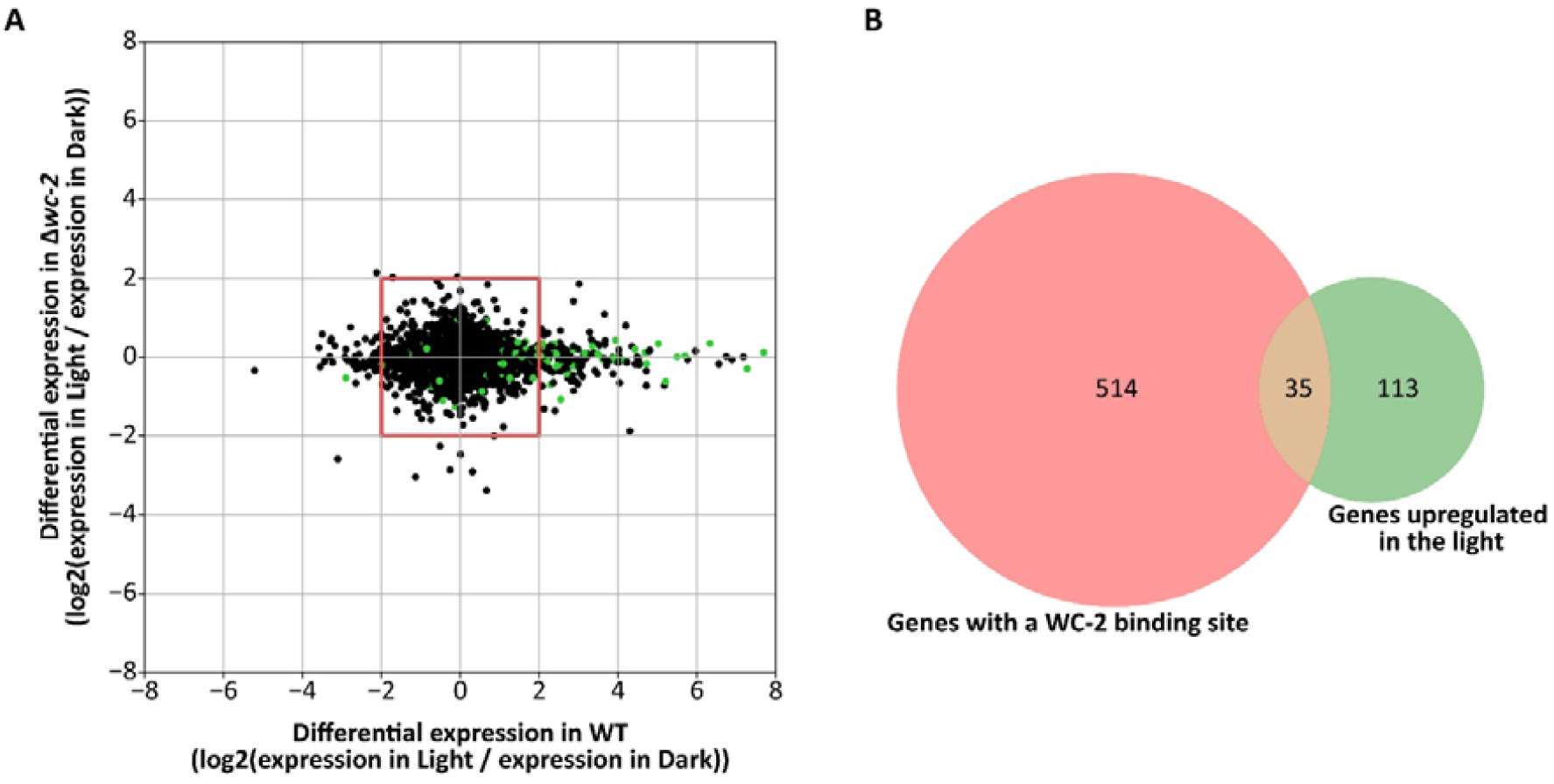
Expression analysis of *S. commune* wild type and Δ*wc-2* dikaryons in the dark and the light. A: Log2 fold-change of all genes in the light compared to the dark in the wild type (x-axis) and Δ*wc-2* dikaryon (y-axis). Genes inside the red box are not considered differentially expressed. Genes highlighted in green are associated with a WC-2 binding site. The majority of genes are only differentially regulated in the WT between light and dark conditions. B: Venn diagram of genes associated with a WC-2 binding site (red) and upregulated DEGs in the WT in the light compared to the dark (green). Many genes presumably regulated by WC-2 are not differentially expressed in the conditions we assayed.

The majority of DEGs in the WT were upregulated in the light (148 out of 195) (Supplementary Table 3). Therefore, light is in general an inducer of expression. It would thus be expected that genes regulated by WC-2 are upregulated in the light in the WT, but not in the Δ*wc-2* strain. This is true for all 148 upregulated DEGs in the WT in the light. These genes are enriched in hydrophobins, oxidoreductases, fatty acid desaturases and a family of genes that encode proteins similar to Bacillus haemolytic enterotoxins (Table 3). Only a single transcription factor, *zfc7*, was differentially upregulated in the light. Of the 148 upregulated DEGs in the light, 35 had a putative WC-2 binding site in the promoter (Figure 4). Of the 47 DEGs upregulated in the dark a single gene was associated with a WC-2 binding site. This indicates that WC-2 is predominantly a positive regulator of transcription and that light alone is not always sufficient for transcription.

**Table 3:**
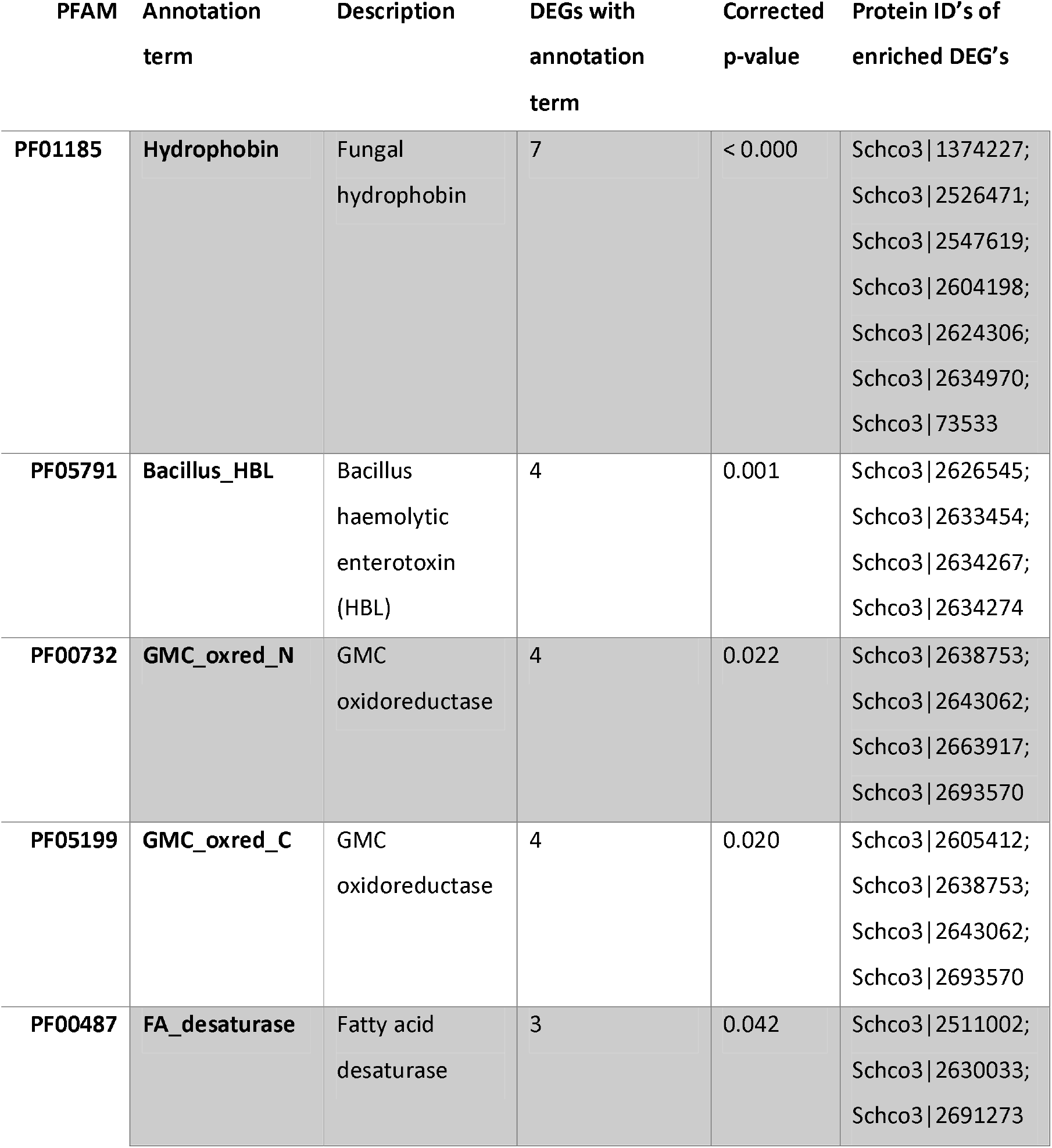
Enrichment analysis of differentially upregulated genes in the light in the wild type. Determined by Fisher’s exact test. P-values are corrected for false discovery rate.

| PFAM | Annotation term | Description | DEGs with annotation term | Corrected p-value | Protein ID's of enriched DEG's |
| --- | --- | --- | --- | --- | --- |
| PF01185 | Hydrophobin | Fungal hydrophobin | 7 | < 0.000 | Schco3\|1374227;<br>Schco3\|2526471;<br>Schco3\|2547619;<br>Schco3\|2604198;<br>Schco3\|2624306;<br>Schco3\|2634970;<br>Schco3\|73533 |
| PF05791 | Bacillus_HBL | Bacillus haemolytic enterotoxin (HBL) | 4 | 0.001 | Schco3\|2626545;<br>Schco3\|2633454;<br>Schco3\|2634267;<br>Schco3\|2634274 |
| PF00732 | GMC_oxred_N | GMC oxidoreductase | 4 | 0.022 | Schco3\|2638753;<br>Schco3\|2643062;<br>Schco3\|2663917;<br>Schco3\|2693570 |
| PF05199 | GMC_oxred_C | GMC oxidoreductase | 4 | 0.020 | Schco3\|2605412;<br>Schco3\|2638753;<br>Schco3\|2643062;<br>Schco3\|2693570 |
| PF00487 | FA_desaturase | Fatty acid desaturase | 3 | 0.042 | Schco3\|2511002;<br>Schco3\|2630033;<br>Schco3\|2691273 |

## Discussion

Light is an important developmental signal in fungi (Corrochano, 2019). In many mushroom-forming fungi, blue light is a key signal for the initiation of fruiting-body development (Morimoto and Oda, 1973; Perkins, 1969; Perkins and Gordon, 1969; Tsusué, 1969). Blue light is detected by the WCC and its downstream gene regulation is essential for the development of mushrooms in *S. commune* (Ohm et al., 2013). Here we describe for the first time the composition of the regulatory network in a mushroom-forming basidiomycete, by a combination of functional genomics, gene expression analysis and transcription factor ChIP-Seq. This reveals that the heterodimerization of WC-1 and WC-2 to form the WCC occurs independently of light. Furthermore, we show that light is necessary for the WCC to associate with the promoters of genes, but that this interaction is not always sufficient to initiate transcription.

As previously described in multiple ascomycetes and the basidiomycete *P. ostreatus*, WC-1 and WC-2 heterodimerize to form the WCC (Corrochano, 2019; Qi et al., 2020). In *S. commune* this action is independent of light, as yeast two-hybrid assays not exposed to blue light still shows interaction of WC-1 and WC-2. This is similar to the previously described heterodimerization of WC-1 and WC-2 in *N. crassa* (Linden and Macino, 1997). In contrast to the *N. crassa* WCC, the complex requires light for interaction with genomic DNA (Sancar et al., 2015). It is tempting to speculate that this is caused by the lack of a zinc-finger DNA-binding domain in the basidiomycete WC-1 ortholog. The presence of two GATA zinc-fingers is often required for the function of these transcription factors (Chen et al., 2012; Hasegawa and Shimizu, 2017). Indeed, many GATA zinc-finger transcription factors contain both an N-terminal and C-terminal domain. For example, in *S. commune* five of the twelve predicted GATA zinc-finger transcription factors contain two GATA zinc-finger domains (Marian et al., 2022). Therefore, a single WCC in ascomycetes may be able to interact with the chromatin independently of light by the dual activity of the WC-1 and WC-2 GATA zinc-finger domains, while light-dependent homodimerization of two WCCs is required in basidiomycetes lacking the WC-1 GATA zinc-finger domain.

A total of 569 WC-2 binding sites, associated with 549 genes were identified for WC-2 during early dikaryotic development. This would indicate that 3.4% of all genes are directly regulated by WC-2, similar to the number of light-regulated genes in ascomycetes (Chen et al., 2009; Idnurm and Heitman, 2010). The consensus motif CGATSNNNNSC was identified in the center of WC-2 binding sites. This motif is very similar to the motif identified for WC-1 that also contains a central CGAT as the primary component of the binding site (Froehlich et al., 2002). Therefore, it appears that the loss of the WC-1 GATA zinc-finger in basidiomycetes does not change the specificity of the WCC.

Despite the large number of putative WC-2 binding sites, only a small subset of 35 of these genes was differentially regulated in the same conditions, indicating that association of WC-2 when exposed to light is not always sufficient for the activation of transcription. This is not surprising, as light is a ubiquitous signal that is known to affect many different processes in fungi. Therefore, other regulatory mechanism may modulate the response to light in addition to WCC. The primary regulator of the WCC in *N. crassa*, VVD, is not found in basidiomycetes (Idnurm et al., 2010; Marian et al., 2022). Nor is there an evident entrainment of a circadian rhythm in *S. commune*. Nevertheless, the overlap in DEGs and genes associated with a WC-2 binding site, indicates that WC-2 has a role in the formation of primordia at the stage we assayed. Multiple hydrophobins that are associated with the structure of primordia, such as *sc1* and *sc4*, are upregulated and associated with WC-2 binding. Moreover, the transcription factor gene *zfc7* was identified in both datasets. This C2H2 zinc-finger was previously shown to have a role in the progression of primordia development (Vonk and Ohm, 2021). This indicates that this process is dependent on light-activation of the WCC, thus promoting the expression of *zfc7*. Therefore, we propose that WC-2 in *S. commune* is a regulator of multiple processes, including early primordia development, but that the role of WC-2 can vary based on other developmental cues and environmental conditions. Previously, two transcription factors with a similar developmental phenotypes as a Δ*wc-2* dikaryon, Hom2 and Tea1, have been identified (Ohm et al., 2011; Pelkmans et al., 2017). WC-2 is associated with the promoters of both of these genes. It is tempting to speculate that these transcription factors work in a cooperative fashion to promote mushroom-development and that this ensures their co-regulation during the initiation of fruiting.

Based on these results and the orthology of the complex between *N. crassa* and *S. commune*, we propose the following mechanism of light detection and subsequent gene regulation in *S. commune* (Figure 5). Both *wc-1* and *wc-2* are constitutively expressed and the translated proteins dimerize to form the WCC. The WCC is not able to associate with chromatin due to the lack of a GATA zinc-finger dimer and is in an inactive state. When light is detected, two WCC homodimerize by the interaction of the LOV domains of WC-1. This homodimerization brings together two GATA zinc-finger domains that bind to the promoters of light-regulated genes to promote transcription. For some genes, this association is sufficient for the initiation of transcription (i.e. photolyase and ferrochelatase), while for others, the recruitment of additional regulators is required to initiate transcription. The nature of these regulators is not yet known, but candidates include additional transcription factors like Hom2 and Tea1. Studies on different time-points of gene regulation are required to identify the full extent of the regulatory network downstream of the detection of light. Furthermore, protein-interaction studies may reveal putative co-factors of the WCC during different environmental and developmental conditions.

**Figure 5:**
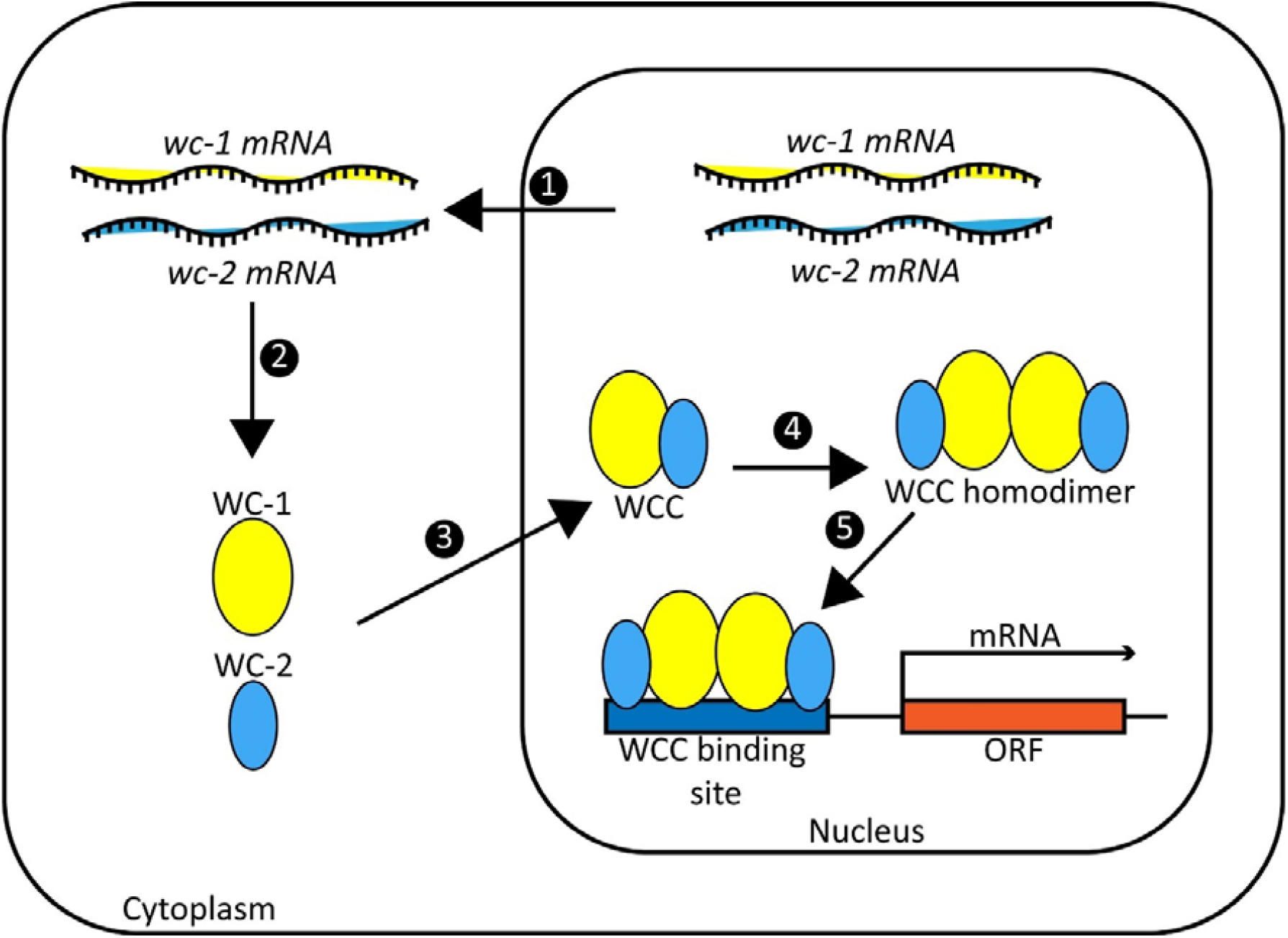
Model for the regulation of transcription by the White Collar complex (WCC) in *S. commune*. 1: Both *wc-1* and *wc-2* are constitutively expressed. 2: mRNA of *wc-1* and *wc-2* is translated into the proteins WC-1 and WC-2. 3: WC-1 and WC-2 dimerize to form the WCC and are translocated to the nucleus due to the presence of a nuclear localization signal on both proteins 4: In the presence of blue light the WCC may dimerize by the interaction of two WC-1 proteins. 5: After dimerization of the WCC, it may become active to promote transcription of light-regulated genes. Co-factors that influence the activity of the WCC have not yet been identified in *S. commune*.

## Materials and Methods

### Strains and culture conditions

The *S. cerevisiae* strains Y8800 (*MATa*) and Y8930 (*MATα*) were a gift from Mike Boxem. These strains with the genotype *trp1-901 leu2-3,112 ura3-52 his3-200 Δgal4 Δgal80 cyh2R GAL1::HIS3@LYS2 GAL2::ADE2 GAL7::LacZ@met2* were routinely grown at 30 °C in yeast extract peptone dextrose medium (YPD), with or without 2% agar for static and liquid cultures respectively. For auxotrophic selection, strains were grown on synthetic complete dropout medium (1.3 g L^-1^ amino acid powder, 1.7 g L^-1^ yeast nitrogen base (Sigma, US), 5 g L^-1^ ammonium sulfate, 2% glucose, pH adjusted to 5.9 with 6M HCl (amino acid powder was 3 g of each alanine, arginine, aspartic acid, asparagine, cysteine, glutamic acid, glutamine, glycine, isoleucine, lysine, methionine, phenylalanine, proline, serine, threonine, tyrosine, uracil, valine)). The medium was supplemented with 1 mM histidine, 1 mM leucine, 0.4 mM tryptophan, 1.68 mM alanine and 0.2 mM 3-amino-1,2,4-triazole depending on the selection conditions.

*S. commune* was grown on *Schizophyllum commune* minimal medium supplemented with 1.5% agar at 25 °C and 30 °C as dikaryons and monokaryons respectively from small agar inoculum (Peer et al., 2009). When grown at 25 °C all cultures were exposed to a 16/8 hours day/night cycle unless specified otherwise. All strains used in this study are derived from *S. commune* H4-8 (*mat*A43*mat*B41; FGSC 9210) (Ohm et al., 2010). For dikaryotic cultures the isogenic compatible strain H4-8b (*mat*A41*matB*43) was used. The *S. commune* Δ*wc-2* strain was published previously (Ohm et al., 2013). For antibiotic selection the medium was supplemented with 15 μg ml^-1^ nourseothricin (Bio-Connect, The Netherlands) or 25 μg ml^-1^ phleomycin (Bio-Connect, The Netherlands).

### Yeast two-hybrid

#### Plasmid construction

Total RNA was isolated with TRIzol (ThermoFisher Scientific, USA) from monokaryotic and dikaryotic H4-8 mycelium grown for 3 days, according to manufacturer’s instructions. Next, this was reverse transcribed to cDNA using the Quantitect Reverse Transcription Kit (Qiagen, Germany). To obtain a more complete library, cDNA from monokaryotic and dikaryotic mycelium was combined. The *wc-1* coding sequence was amplified from *S. commune* H4-8 cDNA with primers wc-1-y2h-prey-fw and wc-1-y2h-prey-rv (Table 4) that contain homology arms to pMB29 around the NotI digestion site, resulting in a 2616 bp fragment. The *wc-2* coding sequence was amplified from *S. commune* H4-8 cDNA with primers wc-2-y2h-bait-fw and wc-2-y2h-bait-rv (Table 4) that contain homology arms to pMB28 around the NotI digestion site, resulting in a 1152 bp fragment. In both forward primers, a guanine was inserted between the homology arm and the gene specific part of the primer, to ensure the gene was inserted in frame. Plasmids pMB29 and pMB28 (a gift from Mike Boxem (Koorman et al., 2016)) were digested with NotI and the *wc-1* and *wc-2* fragments were cloned into the respective plasmids with NEB HiFi DNA assembly mastermix (NEB, US). This resulted in the prey plasmid wc-1-pMB29 and bait plasmid wc-2-pMB28. In wc-1-pMB29, the *wc-1* coding sequence is inserted in frame downstream of the GAL4 activating domain, while in wc-2-pMB28 the *wc-2* coding sequence is inserted in frame downstream of the GAL4 binding domain. Both fusion genes are under the control of the ADH1 promoter and terminator. Furthermore, the prey and bait plasmids contain a TRP1 and LEU2 gene, respectively, to complement the tryptophan and leucine auxotrophy in the Y2H strains.

**Table 4.**
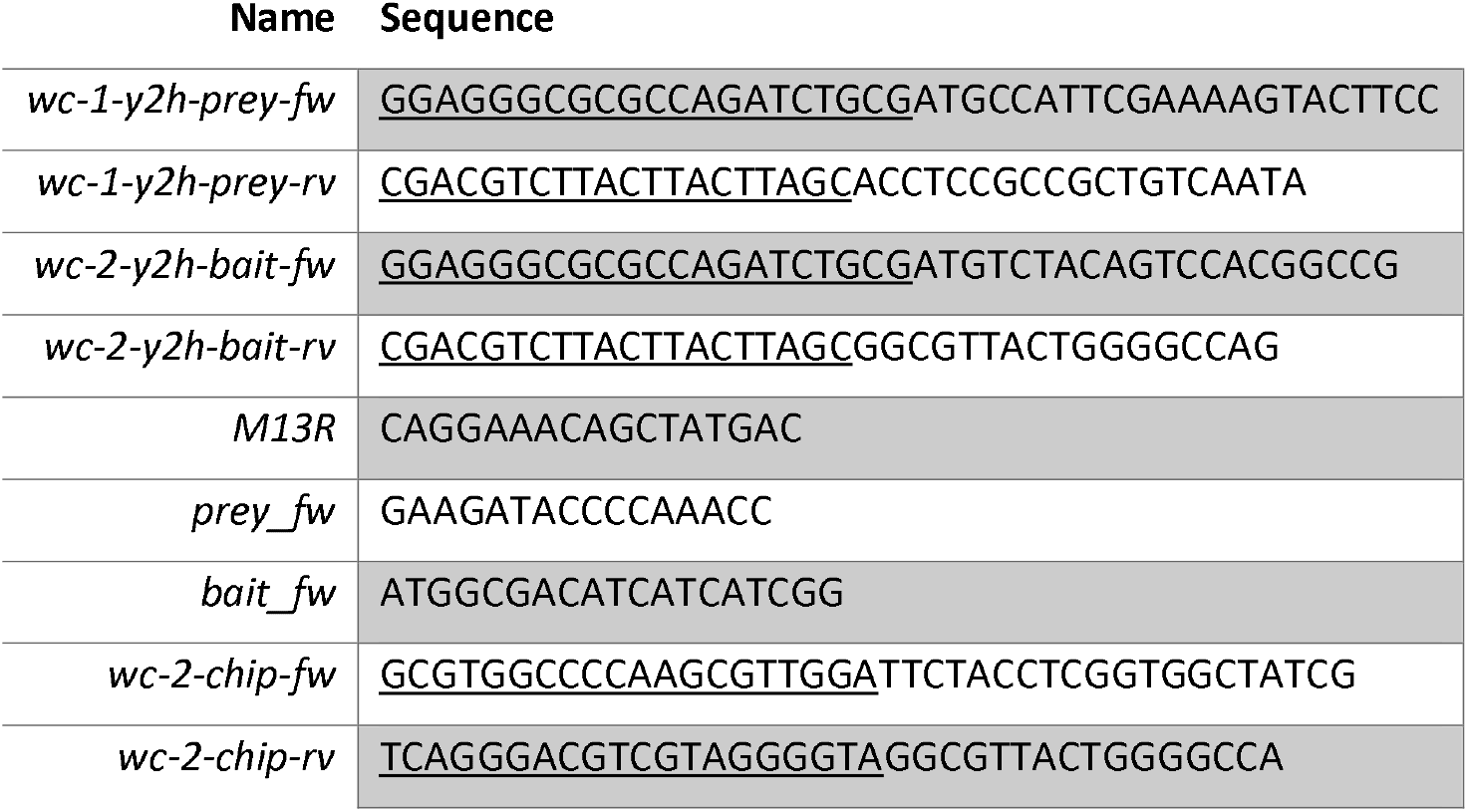
Primers used in this study. Underlined sequences indicate homology arms for Gibson assembly.

#### Yeast transformation and selection

*S. cerevisiae* strains Y8800 and Y8930 were transformed with the plasmids wc-1-pMB29 and wc-2-pMB28, respectively, using the lithium acetate method (Schiestl and Gietz, 1989). Briefly, *S. cerevisiae* was pre-grown overnight at 30 °C at 200 rpm and diluted to an OD_600_ of 0.4 in 50 mL YPD. After growing the culture for an additional 3 hours, the cells were collected by centrifugation at 2500 g for 60 seconds and resuspended in 40 mL 1x TE buffer (10 mM Tris-HCl pH 7.5, 1 mM EDTA pH 8.0). The cells were then centrifuged again at 2500 g for 60 seconds and resuspended in 2 mL LiAc/0.5x TE buffer (100 mM LiAc, 5 mM Tris-HCl pH 7.5, 0.5 mM EDTA pH 8.0). After incubation at room temperature for 10 minutes, 100 μL of the competent cells were mixed with 1 μg plasmid DNA, 100 μg salmon sperm DNA and 700 μL LiAc/PEG-3350/1x TE buffer (100 mM LiAc, 40% w/v PEG-3350, 10 mM Tris-HCl pH 7.5, 1 mM EDTA pH 8.0) and incubated at 30 °C for 30 minutes. Next, 88 μL DMSO was added and cells were heat shocked for 7 minutes at 42 °C. The transformed cells were then collected by brief centrifugation, resuspended in 100 μL 1x TE and plated on synthetic drop-out medium without tryptophan and leucine for strains Y8800 and Y8930, respectively. After incubation at 25 °C for 4 days, transformants were verified with M13R and either prey_fw or bait_fw for preys and baits (Table 4).

#### Mating and yeast two-hybrid

For the yeast two-hybrid the wc-1-pMB29 strain and wc-2-pMB28 strain were mated. As a negative control, strains with empty pMB28 and pMB29 plasmids in Y8930 and Y8800 were used. Each strain was pre-grown overnight in its respective synthetic drop-out medium. Subsequently 10 μL of the respective baits and preys was mixed with 100 μL -Leu -Trp synthetic drop-out medium and centrifuged for 1 minute at 180 rpm. After incubation for 1 day, 15 μL of each cross was transferred to 150 μL -Leu -Trip synthetic drop-out medium and grown for 2 more days. Finally, 15 μL was spotted to a -Leu -Trp -Ade synthetic drop-out medium agar plate and grown for 2 days. For the Y2H experiment without light, mating was performed in monochromatic red light and subsequently incubated in the dark.

### WC-2 ChIP-Seq

#### Plasmid contruction

A plasmid containing the coding sequence of WC-2 fused to an HA-tag under the control of its endogenous promoter was constructed as previously described (Marian et al., 2022). The *wc-2* gene excluding the stop codon and including an 810 bp promoter was amplified from genomic DNA with primers wc-2-chip-fw and wc-2-chip-rv, resulting in a 2165 bp fragment (Table 4). Both primers contained 20 bp homology arms to plasmid pPV009 digested with HindIII. The *wc-2* fragment was cloned into this plasmid with NEB HiFi DNA assembly mastermix creating plasmid wc-2-HA-pPV009.

#### Transformation of S. commune

Plasmid wc-2-HA-pPV009 was transformed into *S. commune* Δ*wc-2* protoplasts as previously described (Peer et al., 2009; Vonk and Ohm, 2021). Successful transformants were selected on phleomycin and confirmed by western blot as previously described (Marian et al., 2022).

#### Chromatin immunoprecipitation

H4-8 and H4-8 Δ*wc-2 :: wc-2-HA* dikaryons were grown on porous polycarbonate (PC) membranes (diameter 76 mm; pore size 0.1 μm; Osmonics; GE Water Technologies, US) for 90 hours in either a 16/8 hour day/night cycle or in the dark. Full colonies were collected and ChIP was performed as previously described in triplicate (Marian et al., 2022). Unlike previously described, ChIP was performed on single colonies of each strain. Furthermore, genomic DNA was fragmented with a diagenode bioruptor UCD-200 (diagenode, US) in 300 μL volume on HIGH setting with a 30 second ON, 30 second OFF cycle for 20 minutes. After DNA isolation libraries were prepared with the NEBNext ULTRA II DNA Library Prep Kit for Illumina (NEB, US) according to manufacturer’s specification with the NEBNext Multiplex Oligos for Illumina (NEB, US). The resulting libraries were sequenced on the Illumina NovaSeq 6000 platform in SP mode with 2x100 bp output at the Utrecht Sequencing Facility (USEQ, www.useq.nl).

#### ChIP-Seq read analysis

ChIP-Seq analysis was performed as previously described (Marian et al., 2022; Vonk and Ohm, 2021). Peaks were called with MACS3 (version 3.0.0b1) with default settings for transcription factor peaks (Zhang et al., 2008). Peaks were then filtered for a minimum score of 100 before manual curation where clear artefacts were removed (i.e., repetitive regions). Peaks were correlated with genes if the center of the peak was within 1000 bp of a translation start site. The distribution of peaks was visualized with deeptools pyGenomeTracks (version 3.8) and custom python scripts (Lopez-Delisle et al., 2021; Ramírez et al., 2018).

#### Motif discovery

Enrichment of motifs was determined with MEME-ChIP from the MEME suite (Bailey et al., 2015). As a positive pool 200 bp fragments around the tip of the ChIP-Seq peaks in WC2-HA in the light were extracted. As a negative control, 10,000 random 200 bp sequences were selected from the genome of *S. commune* H4-8A. The final motif was identified by MEME and the location was determined by Centrimo (both programs are part of MEME suite). The motif was then used in TomTom (part of MEME suite) to search the JASPAR core non-redundant fungal motif database to determine if the motif was previously identified for another transcription factor (Castro-Mondragon et al., 2022).

### RNA Sequencing

H4-8 and H4-8 Δ*wc-2* dikaryons were grown in triplicate on porous polycarbonate (PC) membranes (diameter 76 mm; pore size 0.1 μm; Osmonics; GE Water Technologies, US) for 90 hours in either a 16/8 hour day/night cycle or in the dark. RNA was extracted with TRIzol as previously described. After purification, the RNA was cleaned with the GeneJET RNA Purification kit (ThermoFisher Scientific, USA). Libraries were prepared with the Truseq RNA stranded polyA kit (Illumina, USA) according to manufacturer’s instructions. The resulting libraries were sequenced on the Illumina NextSeq 2000 platform with a P2 flowcell and 2x50 bp output at the Utrecht Sequencing Facility (USEQ, http://www.useq.nl) Reads were aligned with HISAT2 version 2.2.1 (Kim et al., 2019) and differential expression was determined with Cuffdiff version 2.2.1 (Trapnell et al., 2013). Genes were considered differentially regulated when Cuffdiff considered them significant, and there was a 4-fold change in expression between any conditions, and the minimum expression in either condition was > 10 RPKM.

## Data availability

The RNA sequencing and ChIP sequencing reads have been deposited in the NCBI Short Read Archive and can be accessed under bioproject ID PRJNA1073722.

## Acknowledgements

This project has received funding from the European Research Council (ERC) under the European Union’s Horizon 2020 research and innovation program (grant agreement number 716132). We acknowledge the Utrecht Sequencing Facility (USEQ) for providing sequencing service and data. USEQ is subsidized by the University Medical Center Utrecht and The Netherlands X-omics Initiative (NWO project 184.034.019).

## Author contributions

Performed experiments and analyzed the data: PJV, MJPP, ZEN. Supervision/coordination: PJV, RAO. Wrote the manuscript: PJV, RAO. Provided funding: RAO. Designed the experiments: PJV, RAO. Read and approved the manuscript: All Authors.

---

**Supplementary Figure 1:**
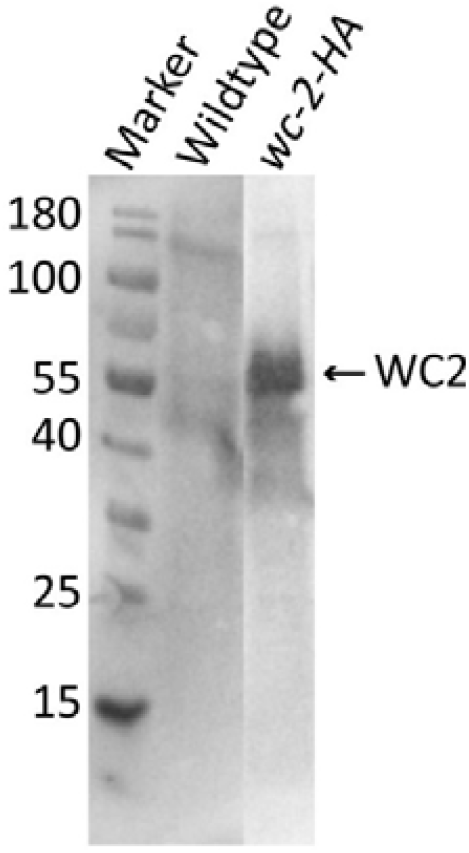
Western blot of WC-2 fused to and HA-tag. The expected band size without any posttranslational modification is 43 kDa.

**Supplementary Figure 2:**
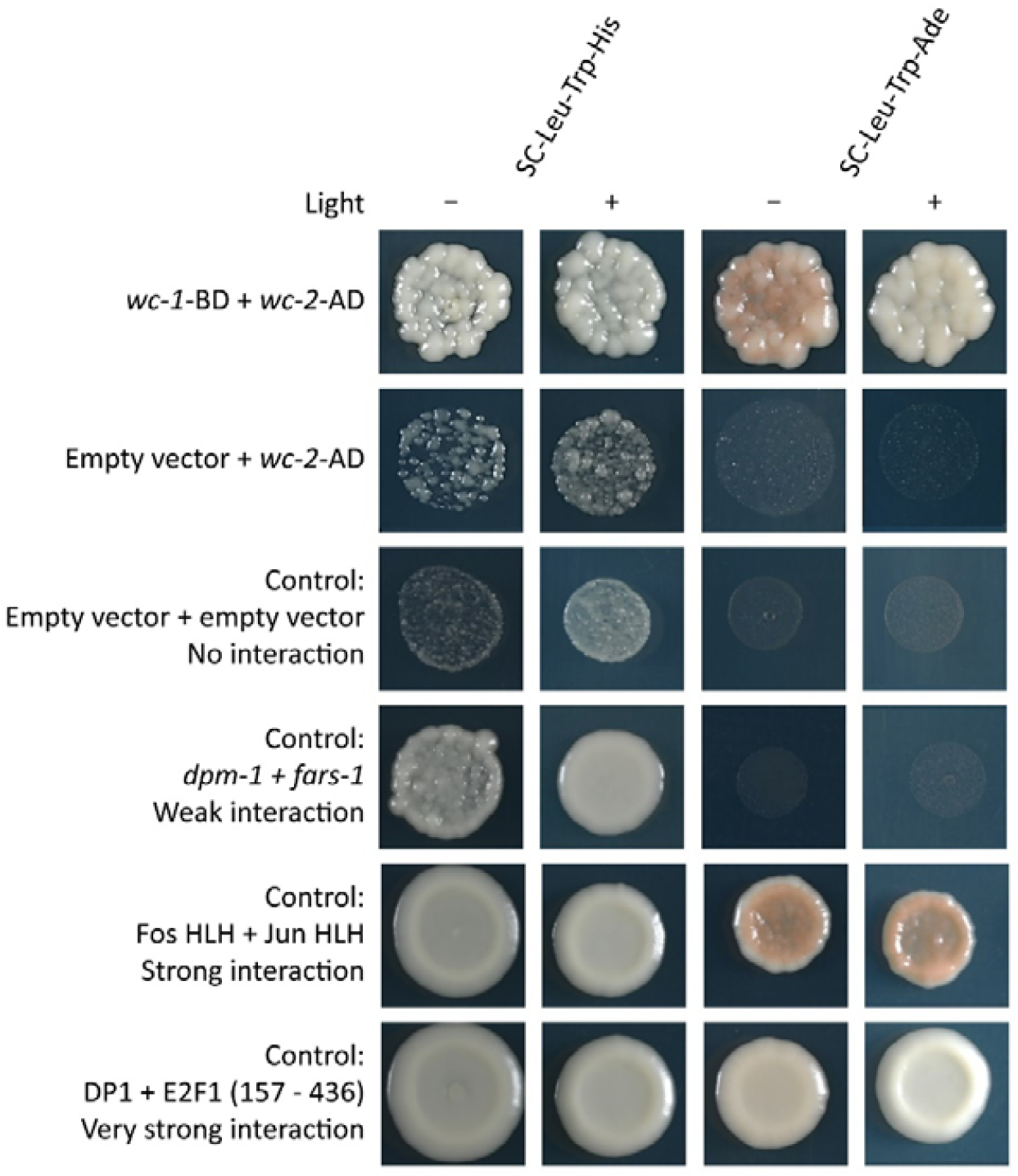
Yeast two-hybrid of *wc-1* fused to the Gal4-activating domain (AD) and *wc-2* fused to the Gal4-binding domain (BD) in the light and the dark. Selection was performed on synthetic medium lacking histidine (-His) and adenine (-Ade). Stronger interaction is required for the complementation of adenine auxotrophy. As a negative control an empty prey vector was used. Furthermore, four controls of varying interaction ranging from no interaction to very strong interaction were used (Boxem et al., 2008).

**Supplementary Table 1: ChIP-Seq peaks of WC-2-HA in the light and the dark as identified by ChIP-Seq. The protein IDs of genes with a start site within 1000 bp of the tip of a peak**.

**Supplementary Table 2: Genes associated with a WC-2 binding site and their functional annotation**.

**Supplementary Table 3: Differentially expressed genes in the wild type and Δ*wc-2* dikaryons grown in the light and dark for 90 hours**.

**Supplementary Table 4: Gene expression values (in RPKM) in the wild type and Δ*wc-2* dikaryons grown in the light and dark for 90 hours**.

## References

Bailey, T.L., Johnson, J., Grant, C.E., Noble, W.S., 2015. The MEME Suite. Nucleic Acids Res. 43, W39–W49. 10.1093/nar/gkv416

Ballario, P., Vittorioso, P., Magrelli, A., Talora, C., Cabibbo, A., Macino, G., 1996. White collar-1, a central regulator of blue light responses in Neurospora, is a zinc finger protein. EMBO J. 15, 1650–1657. 10.1002/j.1460-2075.1996.tb00510.x

Boxem, M., Maliga, Z., Klitgord, N., Li, N., Lemmens, I., Mana, M., Lichtervelde, L. de, Mul, J.D., Peut, D. van de, Devos, M., Simonis, N., Yildirim, M.A., Cokol, M., Kao, H.-L., Smet, A.-S., de Wang, H., Schlaitz, A.-L., Hao, T., Milstein, S., Fan, C., Tipsword, M., Drew, K., Galli, M., Rhrissorrakrai, K., Drechsel, D., Koller, D., Roth, F.P., Iakoucheva, L.M., Dunker, A.K., Bonneau, R., Gunsalus, K.C., Hill, D.E., Piano, F., Tavernier, J., Heuvel, S. van den, Hyman, A.A., Vidal, M., 2008. A Protein Domain-Based Interactome Network for C. elegans Early Embryogenesis. Cell 134, 534–545. 10.1016/j.cell.2008.07.009

Castro-Mondragon, J.A., Riudavets-Puig, R., Rauluseviciute, I., Berhanu Lemma, R., Turchi, L., Blanc-Mathieu, R., Lucas, J., Boddie, P., Khan, A., Manosalva Pérez, N., Fornes, O., Leung, T.Y., Aguirre, A., Hammal, F., Schmelter, D., Baranasic, D., Ballester, B., Sandelin, A., Lenhard, B., Vandepoele, K., Wasserman, W.W., Parcy, F., Mathelier, A., 2022. JASPAR 2022: the 9th release of the open-access database of transcription factor binding profiles. Nucleic Acids Res. 50, D165–D173. 10.1093/nar/gkab1113

Chen, C., Ringelberg, C.S., Gross, R.H., Dunlap, J.C., Loros, J.J., 2009. Genome-wide analysis of light-inducible responses reveals hierarchical light signalling in Neurospora. EMBO J. 28, 1029–1042. 10.1038/emboj.2009.54

Chen, C.-H., Dunlap, J.C., Loros, J.J., 2010. Neurospora illuminates fungal photoreception. Fungal Genet. Biol. FG B 47, 922–929. 10.1016/j.fgb.2010.07.005

Chen, Y., Bates, D.L., Dey, R., Chen, P.-H., Machado, A.C.D., Laird-Offringa, I.A., Rohs, R., Chen, L., 2012. DNA binding by GATA transcription factor suggests mechanisms of DNA looping and long-range gene regulation. Cell Rep. 2, 1197–1206. 10.1016/j.celrep.2012.10.012

Corrochano, L.M., 2019. Light in the Fungal World: From Photoreception to Gene Transcription and Beyond. Annu. Rev. Genet. 53, 149–170. 10.1146/annurev-genet-120417-031415

De Jong, J.F., Ohm, R.A., De Bekker, C., Wösten, H.A.B., Lugones, L.G., 2010. Inactivation of ku80 in the mushroom-forming fungus Schizophyllum commune increases the relative incidence of homologous recombination. FEMS Microbiol. Lett. 310, 91–95. 10.1111/j.1574-6968.2010.02052.x

Dunlap, J.C., Loros, J.J., 2017. Making Time: Conservation of Biological Clocks from Fungi to Animals. Microbiol. Spectr. 5, 5.3.05. 10.1128/microbiolspec.FUNK-0039-2016

Dunlap, J.C., Loros, J.J., 2005. Neurospora Photoreceptors, in: Handbook of Photosensory Receptors. John Wiley & Sons, Ltd, pp. 371–389. 10.1002/352760510X.ch18

Froehlich, A.C., Liu, Y., Loros, J.J., Dunlap, J.C., 2002. White Collar-1, a Circadian Blue Light Photoreceptor, Binding to the frequency Promoter. Science 297, 815–819. 10.1126/science.1073681

Grimaldi, B., Coiro, P., Filetici, P., Berge, E., Dobosy, J.R., Freitag, M., Selker, E.U., Ballario, P., 2006. The Neurospora crassa White Collar-1 dependent Blue Light Response Requires Acetylation of Histone H3 Lysine 14 by NGF-1. Mol. Biol. Cell 17, 4576–4583. 10.1091/mbc.e06-03-0232

Hasegawa, A., Shimizu, R., 2017. GATA1 Activity Governed by Configurations of cis-Acting Elements. Front. Oncol. 6.

He, Q., Cheng, P., Yang, Y., Wang, L., Gardner, K.H., Liu, Y., 2002. White Collar-1, a DNA Binding Transcription Factor and a Light Sensor. Science 297, 840–843. 10.1126/science.1072795

Idnurm, A., Heitman, J., 2010. Ferrochelatase is a conserved downstream target of the blue light-sensing White collar complex in fungi. Microbiology 156, 2393–2407. 10.1099/mic.0.039222-0

Idnurm, A., Verma, S., Corrochano, L.M., 2010. A glimpse into the basis of vision in the kingdom Mycota. Fungal Genet. Biol., Special Issue: Photobiology 47, 881–892. 10.1016/j.fgb.2010.04.009

Kamada, T., Sano, H., Nakazawa, T., Nakahori, K., 2010. Regulation of fruiting body photomorphogenesis in Coprinopsis cinerea. Fungal Genet. Biol., Special Issue: Photobiology 47, 917–921. 10.1016/j.fgb.2010.05.003

Kim, D., Paggi, J.M., Park, C., Bennett, C., Salzberg, S.L., 2019. Graph-based genome alignment and genotyping with HISAT2 and HISAT-genotype. Nat. Biotechnol. 37, 907–915. 10.1038/s41587-019-0201-4

Koorman, T., Klompstra, D., Voet, M. van der, Lemmens, I., Ramalho, J.J., Nieuwenhuize, S., Heuvel, S. van den, Tavernier, J., Nance, J., Boxem, M., 2016. A combined binary interaction and phenotypic map of C. elegans cell polarity proteins. Nat. Cell Biol. 18, 337. 10.1038/ncb3300

Li, J., Xu, C., Jing, Z., Li, X., Li, H., Chen, Y., Shao, Y., Cai, J., Wang, B., Xie, B., Tao, Y., 2023. Blue light and its receptor white collar complex (FfWCC) regulates mycelial growth and fruiting body development in Flammulina filiformis. Sci. Hortic. 309, 111623. 10.1016/j.scienta.2022.111623

Linden, H., Macino, G., 1997. White collar 2, a partner in blue-light signal transduction, controlling expression of light–regulated genes in Neurospora crassa. EMBO J. 16, 98–109. 10.1093/emboj/16.1.98

Lopez-Delisle, L., Rabbani, L., Wolff, J., Bhardwaj, V., Backofen, R., Grüning, B., Ramírez, F., Manke, T., 2021. pyGenomeTracks: reproducible plots for multivariate genomic datasets. Bioinformatics 37, 422–423. 10.1093/bioinformatics/btaa692

Malzahn, E., Ciprianidis, S., Káldi, K., Schafmeier, T., Brunner, M., 2010. Photoadaptation in Neurospora by Competitive Interaction of Activating and Inhibitory LOV Domains. Cell 142, 762–772. 10.1016/j.cell.2010.08.010

Marian, I.M., Vonk, P.J., Valdes, I.D., Barry, K., Bostock, B., Carver, A., Daum, C., Lerner, H., Lipzen, A., Park, H., Schuller, M.B.P., Tegelaar, M., Tritt, A., Schmutz, J., Grimwood, J., Lugones, L.G., Choi, I.-G., Wösten, H.A.B., Grigoriev, I.V., Ohm, R.A., 2022. The Transcription Factor Roc1 Is a Key Regulator of Cellulose Degradation in the Wood-Decaying Mushroom Schizophyllum commune. mBio 13, e00628–22. 10.1128/mbio.00628-22

Morimoto, N., Oda, Y., 1973. Effects of light on fruit-body formation in a basidiomycete, Coprinus macrorhizus. Plant Cell Physiol. 14, 217–225. 10.1093/oxfordjournals.pcp.a074854

Ohm, R.A., Aerts, D., Wösten, H.A.B., Lugones, L.G., 2013. The blue light receptor complex WC-1/2 of Schizophyllum commune is involved in mushroom formation and protection against phototoxicity. Environ. Microbiol. 15, 943–955. 10.1111/j.1462-2920.2012.02878.x

Ohm, R.A., de Jong, J.F., de Bekker, C., Wösten, H.A.B., Lugones, L.G., 2011. Transcription factor genes of Schizophyllum commune involved in regulation of mushroom formation. Mol. Microbiol. 81, 1433–1445. 10.1111/j.1365-2958.2011.07776.x

Ohm, R.A., de Jong, J.F., Lugones, L.G., Aerts, A., Kothe, E., Stajich, J.E., de Vries, R.P., Record, E., Levasseur, A., Baker, S.E., Bartholomew, K.A., Coutinho, P.M., Erdmann, S., Fowler, T.J., Gathman, A.C., Lombard, V., Henrissat, B., Knabe, N., Kües, U., Lilly, W.W., Lindquist, E., Lucas, S., Magnuson, J.K., Piumi, F., Raudaskoski, M., Salamov, A., Schmutz, J., Schwarze, F.W.M.R., vanKuyk, P.A., Horton, J.S., Grigoriev, I.V., Wösten, H.A.B., 2010. Genome sequence of the model mushroom Schizophyllum commune. Nat. Biotechnol. 28, 957–963. 10.1038/nbt.1643

Peer, A.F. van Müller, W.H., Boekhout, T., Lugones, L.G., Wösten, H.A.B., 2009. Cytoplasmic Continuity Revisited: Closure of Septa of the Filamentous Fungus Schizophyllum commune in Response to Environmental Conditions. PLoS ONE 4. 10.1371/journal.pone.0005977

Pelkmans, J.F., Patil, M.B., Gehrmann, T., Reinders, M.J.T., Wösten, H.A.B., Lugones, L.G., 2017. Transcription factors of Schizophyllum commune involved in mushroom formation and modulation of vegetative growth. Sci. Rep. 7, 310. 10.1038/s41598-017-00483-3

Perkins, J.H., 1969. Morphogenesis in Schizophyllum commune. I. Effects of White Light 1. Plant Physiol. 44, 1706–1711. 10.1104/pp.44.12.1706

Perkins, J.H., Gordon, S.A., 1969. Morphogenesis in Schizophyllum commune. II. Effects of Monochromatic Light 1. Plant Physiol. 44, 1712–1716. 10.1104/pp.44.12.1712

Qi, Y., Sun, X., Ma, L., Wen, Q., Qiu, L., Shen, J., 2020. Identification of two Pleurotus ostreatus blue light receptor genes (PoWC-1 and PoWC-2) and in vivo confirmation of complex PoWC-12 formation through yeast two hybrid system. Fungal Biol. 124, 8–14. 10.1016/j.funbio.2019.10.004

Ramírez, F., Bhardwaj, V., Arrigoni, L., Lam, K.C., Grüning, B.A., Villaveces, J., Habermann, B., Akhtar, A., Manke, T., 2018. High-resolution TADs reveal DNA sequences underlying genome organization in flies. Nat. Commun. 9, 189. 10.1038/s41467-017-02525-w

Sancar, C., Ha, N., Yilmaz, R., Tesorero, R., Fisher, T., Brunner, M., Sancar, G., 2015. Combinatorial Control of Light Induced Chromatin Remodeling and Gene Activation in Neurospora. PLOS Genet. 11, e1005105. 10.1371/journal.pgen.1005105

Schafmeier, T., Haase, A., Káldi, K., Scholz, J., Fuchs, M., Brunner, M., 2005. Transcriptional Feedback of Neurospora Circadian Clock Gene by Phosphorylation-Dependent Inactivation of Its Transcription Factor. Cell 122, 235–246. 10.1016/j.cell.2005.05.032

Schiestl, R.H., Gietz, R.D., 1989. High efficiency transformation of intact yeast cells using single stranded nucleic acids as a carrier. Curr. Genet. 16, 339–346. 10.1007/BF00340712

Todd, R.B., Zhou, M., Ohm, R.A., Leeggangers, H.A., Visser, L., de Vries, R.P., 2014. Prevalence of transcription factors in ascomycete and basidiomycete fungi. BMC Genomics 15, 214. 10.1186/1471-2164-15-214

Trapnell, C., Hendrickson, D.G., Sauvageau, M., Goff, L., Rinn, J.L., Pachter, L., 2013. Differential analysis of gene regulation at transcript resolution with RNA-seq. Nat. Biotechnol. 31, 46–53. 10.1038/nbt.2450

Tsusué, Y.M., 1969. Experimental Control of Fruit-Body Formation in Coprinus Macrorhizus. Dev. Growth Differ. 11, 164–178. 10.1111/j.1440-169X.1969.00164.x

Vonk, P.J., Ohm, R.A., 2021. H3K4me2 ChIP-Seq reveals the epigenetic landscape during mushroom formation and novel developmental regulators of Schizophyllum commune. Sci. Rep. 11, 8178. 10.1038/s41598-021-87635-8

Zhang, Y., Liu, T., Meyer, C.A., Eeckhoute, J., Johnson, D.S., Bernstein, B.E., Nusbaum, C., Myers, R.M., Brown, M., Li, W., Liu, X.S., 2008. Model-based Analysis of ChIP-Seq (MACS). Genome Biol. 9, R137. 10.1186/gb-2008-9-9-r137

